# Platelet-Localized ST6Gal1 Does Not Impact IgG Sialylation

**DOI:** 10.1101/2023.06.15.545133

**Authors:** Leandre M. Glendenning, Julie Y. Zhou, Emily N. Kukan, Chao Gao, Richard D. Cummings, Smita Joshi, Sidney W. Whiteheart, Brian A. Cobb

**Affiliations:** Department of Pathology, Case Western Reserve University School of Medicine; Harvard Medical School Center for Glycoscience, National Center for Functional Glycomics, Beth Israel Deaconess Medical Center; Department of Molecular and Cellular Biochemistry, University of Kentucky

**Keywords:** platelet, IgG, sialylation, ST6Gal1, sialic acid

## Abstract

The IgG antibody class forms an important basis of the humoral immune response, conferring reciprocal protection from both pathogens and autoimmunity. IgG function is determined by the IgG subclass, as defined by the heavy chain, as well as the glycan composition at N297, the conserved site of N-glycosylation within the Fc domain. For example, lack of core fucose promotes increased antibody-dependent cellular cytotoxicity, whereas α2,6-linked sialylation by the enzyme ST6Gal1 helps to drive immune quiescence. Despite the immunological significance of these carbohydrates, little is known about how IgG glycan composition is regulated. We previously reported that mice with ST6Gal1-deficient B cells have unaltered IgG sialylation. Likewise, ST6Gal1 released into the plasma by hepatocytes does not significantly impact overall IgG sialylation. Since IgG and ST6Gal1 have independently been shown to exist in platelet granules, it was possible that platelet granules could serve as a B cell-extrinsic site for IgG sialylation. To address this hypothesis, we used a platelet factor 4 (Pf4)-Cre mouse to delete ST6Gal1 in megakaryocytes and platelets alone or in combination with an albumin-Cre mouse to also remove it from hepatocytes and the plasma. The resulting mouse strains were viable and had no overt pathological phenotype. We also found that despite targeted ablation of ST6Gal1, no change in IgG sialylation was apparent. Together with our prior findings, we can conclude that in mice, neither B cells, the plasma, nor platelets have a substantial role in homeostatic IgG sialylation.

## Introduction

IgG is one of the most abundant proteins in the plasma and performs a variety of immunomodulatory functions, which are primarily driven by interactions between the Fc domain and a variety of cell surface receptors. The Fc domain contains a single conserved N-glycosylation site at Asn297, and the glycans at this site are relatively conserved in structure, nearly always being biantennary with variations in composition, such as the presence or absence of bisecting GlcNAc, core fucose, galactose, and sialic acid (de Haan, N., Reiding, K.R., et al. 2017). Interestingly, the composition of this glycan, particularly the presence or absence of the terminal α2,6-linked sialic acid and its underlying galactose, have been associated with a variety of autoimmune and infectious diseases (Cobb, B.A. 2020). Early studies demonstrated an association between asialylated IgG and symptomatic rheumatoid arthritis (RA), which often enters remission during pregnancy along with a concomitant increase in plasma IgG sialylation (Pekelharing, J.M., Hepp, E., et al. 1988). These studies show that IgG sialylation is dynamically regulated, changing in concordance with the overall organismal state. Increases in IgG sialylation increase anti-inflammatory effector functions, often by enhancing binding to the inhibitory Fc receptor FcγRIIb, and enriching IVIg preparations for sialylated species even increases the therapeutic efficacy of this anti-inflammatory therapy (Kaneko, Y., Nimmerjahn, F., et al. 2006, Schwab, I., Biburger, M., et al. 2012). More recently, changes in IgG glycan composition as a result of specific vaccine adjuvants have been identified (Mahan, A.E., Jennewein, M.F., et al. 2016), suggesting that the composition of the IgG glycan is tightly regulated in a manner specific to the inflammatory insult and that a currently unidentified mechanism integrates the detection of inflammatory mediators with pathways underlying IgG sialylation.

N-linked glycans are added to a nascent protein during synthesis in the rough endoplasmic reticulum and are modified within the Golgi apparatus of the secretory pathway, with individual monosaccharides added to the glycan in a roughly sequential manner. Sialic acids, as a terminal modification, are added to the glycan of IgG in the *trans-*Golgi network. It is currently believed that glycoprotein glycans can only be modified in composition as they are being secreted, and thus it is assumed that B cells modulate IgG sialylation. However, we previously published that a mouse strain selectively lacking ST6Gal1 in B cells possessed normal amounts of sialylated IgG in the plasma (Jones, M.B., Oswald, D.M., et al. 2016). More recently, we showed that B cells secrete predominantly asialylated IgG into the bloodstream as a result of unique intracellular protein trafficking pathways that significantly reduce IgG contact with ST6Gal1 inside of the cell (Glendenning, L.M., Zhou, J.Y., et al. 2022). In combination with previously published literature on the dynamic nature of IgG sialylation in response to various inflammatory conditions (Cobb, B.A. 2020), this suggests a model in which B cells secrete asialylated IgG into the bloodstream as a default, and that sialylation occurs within a different compartment in accordance with the current immunologic state.

As approximately 15% of serum IgG is sialylated (Gudelj, I., Lauc, G., et al. 2018), we first hypothesized that IgG sialylation was occurring in the plasma as a result of circulating ST6Gal1. Supporting this hypothesis, circulating enzyme secreted by the liver was shown to be active in the circulation, and knocking hepatocyte-derived ST6Gal1 out of mice ablated the sialylation status of the vast majority of plasma glycoproteins (Oswald, D.M., Lehoux, S.D., et al. 2022). In addition, ablation of the P1 promoter of ST6Gal1, which is reportedly liver-specific (Dalziel, M., Lemaire, S., et al. 1999), results in a reduction of IgG sialylation (Jones, M.B., Nasirikenari, M., et al. 2012). Despite nearly all of the other plasma glycoproteins being impacted by the lack of hepatocyte-derived enzyme, the IgG in the hepatocyte-specific ST6Gal1 knockout animal remained normally sialylated, suggesting that IgG is not sialylated directly in the plasma (Oswald, D.M., Lehoux, S.D., et al. 2022).

These findings lead to the general hypothesis that IgG must first be endocytosed by another cell, placing the protein within a potential *in vivo* reaction vessel that includes the sialyltransferase ST6Gal1, IgG, and the sugar donor CMP-sialic acid to enable efficient sialylation. As endogenous IgG and exogenous IgG-based drugs are known to be endocytosed into platelets and even used as a marker for α-granules (George, J.N. 1991, Verheul, H.M., Lolkema, M.P., et al. 2007), we hypothesized that these granules may be this elusive intracellular location of IgG sialylation. This notion was supported by our data showing that mouse platelets contain the sugar donor CMP-sialic acid, and when activated, can release a sufficient quantity of CMP-sialic acid to sialylate IgG *in vitro* (Jones, M.B., Oswald, D.M., et al. 2016). Additionally, ST6Gal1 and CMP-SA have been quantified within human platelet granules (Wandall, H.H., Rumjantseva, V., et al. 2012), potentially allowing for endocytosed IgG to be sialylated and ultimately released upon platelet degranulation. Therefore, we generated a novel murine strain that lacks ST6Gal1 in platelets (PcKO). Here, we describe the impact of this knockout on the phenotype of the animal as well as its impact on IgG sialylation. Our results indicate that IgG sialylation remains unchanged despite the lack of ST6Gal1 expression in platelets. Further supporting this conclusion, we found that other murine strains with platelet secretion deficiencies did not result in the decrease of IgG sialylation. To rule out the possibility that platelet- and hepatocyte-derived (i.e. plasma-localized) ST6Gal1 are able to functionally compensate for the loss of the enzyme in either cell, we further developed a murine strain lacking ST6Gal1 in both platelets and hepatocytes. The sialylation status of IgG in this animal remained unchanged, demonstrating that platelets do not serve as the reaction vessel for B cell-extrinsic IgG sialylation.

## Results

### Lack of ST6Gal1 in platelets with and without circulatory ST6Gal1 generates modest non-pathogenic changes

Pf4-Cre and Albumin-Cre mice were used in combination with a strain carrying ST6Gal1-flanking LoxP sites to enable selective ablation of the enzyme in megakaryocytes (MKs) and platelets (Fig 1A) as well as in hepatocytes (Fig 1B), respectively. When crossed, we therefore generated two novel mouse strains: a MK/platelet-specific conditional knockout of ST6Gal1 (PcKO) with reduced α2,6-sialylation on platelets (Fig 1C) and a combined platelet and hepatocyte-specific conditional knockout (PHcKO). Since platelets endocytose plasma components, including IgG and potentially ST6Gal1, it was critical to remove ST6Gal1 from the hepatocytes, which eliminates circulatory ST6Gal1 and thus any possibility that transferase activity could remain within platelet granules (Oswald, D.M., Lehoux, S.D., et al. 2022). As we have previously reported, the hepatocyte-specific conditional knockout of ST6Gal1 (HcKO) results in a loss of α2,6-sialylation on the surface of hepatocytes and on most plasma-localized glycoproteins (Oswald, D.M., Jones, M.B., et al. 2020), excluding IgG (Oswald, D.M., Lehoux, S.D., et al. 2022). The HcKO model also does not show overt pathology in young animals, exhibiting no changes in blood chemistry or cellularity, but develops a progressive liver disease by approximately 40 weeks (Oswald, D.M., Jones, M.B., et al. 2020).

**Figure 1.**
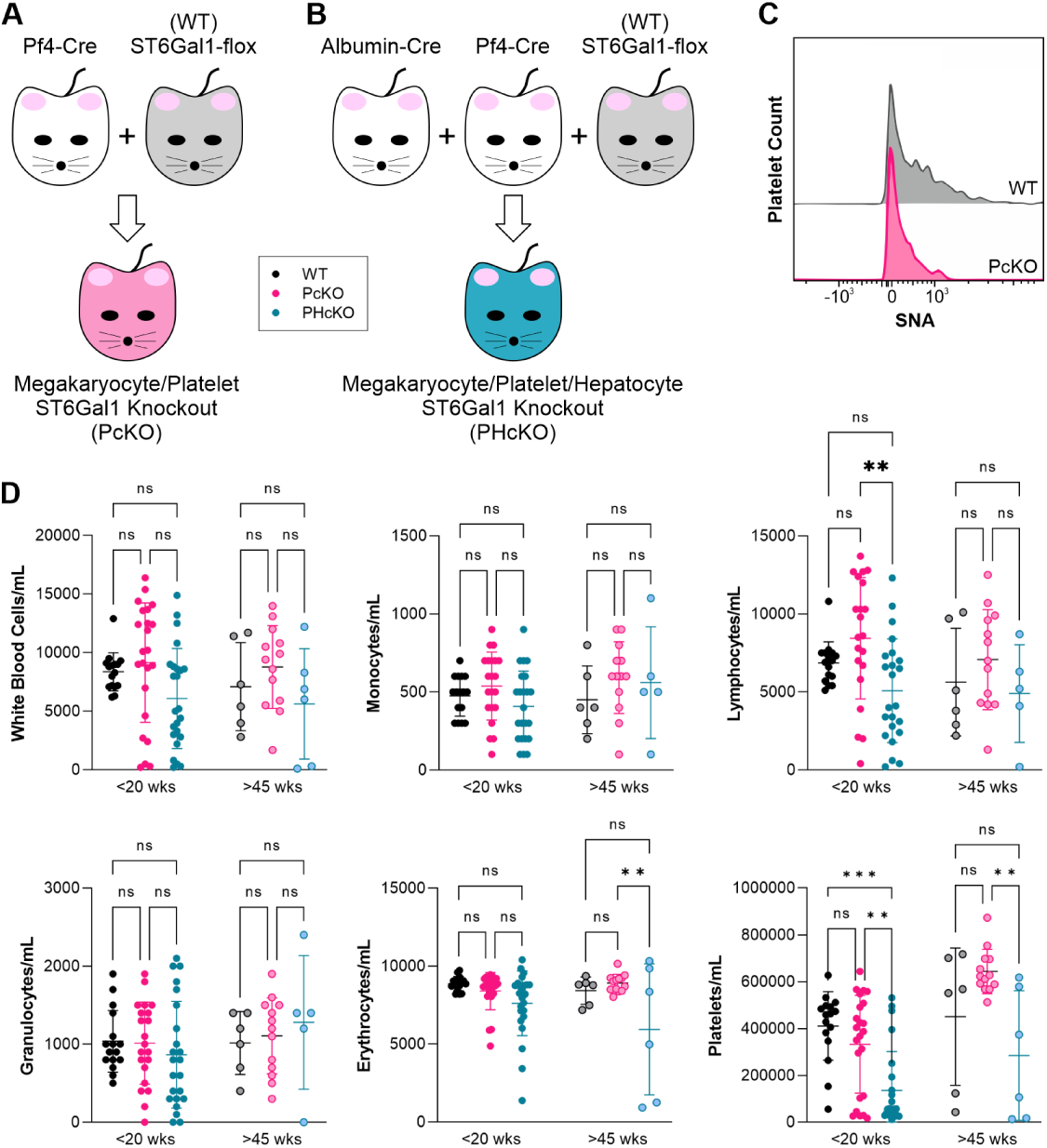
Creation and characterization of blood cellularity of the PcKO and PHcKO mice. (A) Scheme for the generation of the MK/platelet-specific ablation of ST6Gal1 in mice. (B) Scheme for the generation of the dual MK/platelet-hepatocyte-specific ablation of ST6Gal1 in mice. (C) Flow cytometry of CD31+ platelets from WT and PcKO animals stained with SNA, showing marked reduction in cell surface α2,6-linked sialic acids. (D) Blood cellular differentials from WT, PcKO, and PHcKO animals grouped by age. Data represents both male and female animals. **p < 0.01; ***p < 0.001; N=6-24

In order to characterize the broad impact of ST6Gal1 ablation in PcKO and PHcKO animals, we first compared the blood cellularity of young animals less than 20 weeks of age and old animals greater than 45 weeks of age to the ST6Gal1-flox WT animals. We found that most cell counts were indistinguishable from WT within both age groups (Fig 1D). However, we did find that lymphocyte numbers were widely distributed, particularly in both mutant strains, with the young PHcKO mice showing a significant reduction compared to young WT animals. We found a similarly wide distribution of erythrocytes in old PHcKO mice and an overall reduction in the number of platelets in the PHcKO mouse compared to either WT or PcKO animals (Fig 1D). These findings suggest that circulatory ST6Gal1 activity impacts platelets directly, although the mechanism for this observation has yet to be investigated.

Next, we performed a survey of major tissues by H&E staining (Fig 2). We found a lack of any notable pathologic change in any tissue within either age group, including the liver from aged PHcKO mice which showed only very minor changes in tissue architecture. Importantly, the number and organization of splenic germinal centers was not impacted by the genetic alterations, suggesting that IgG responses should not be altered.

**Figure 2.**
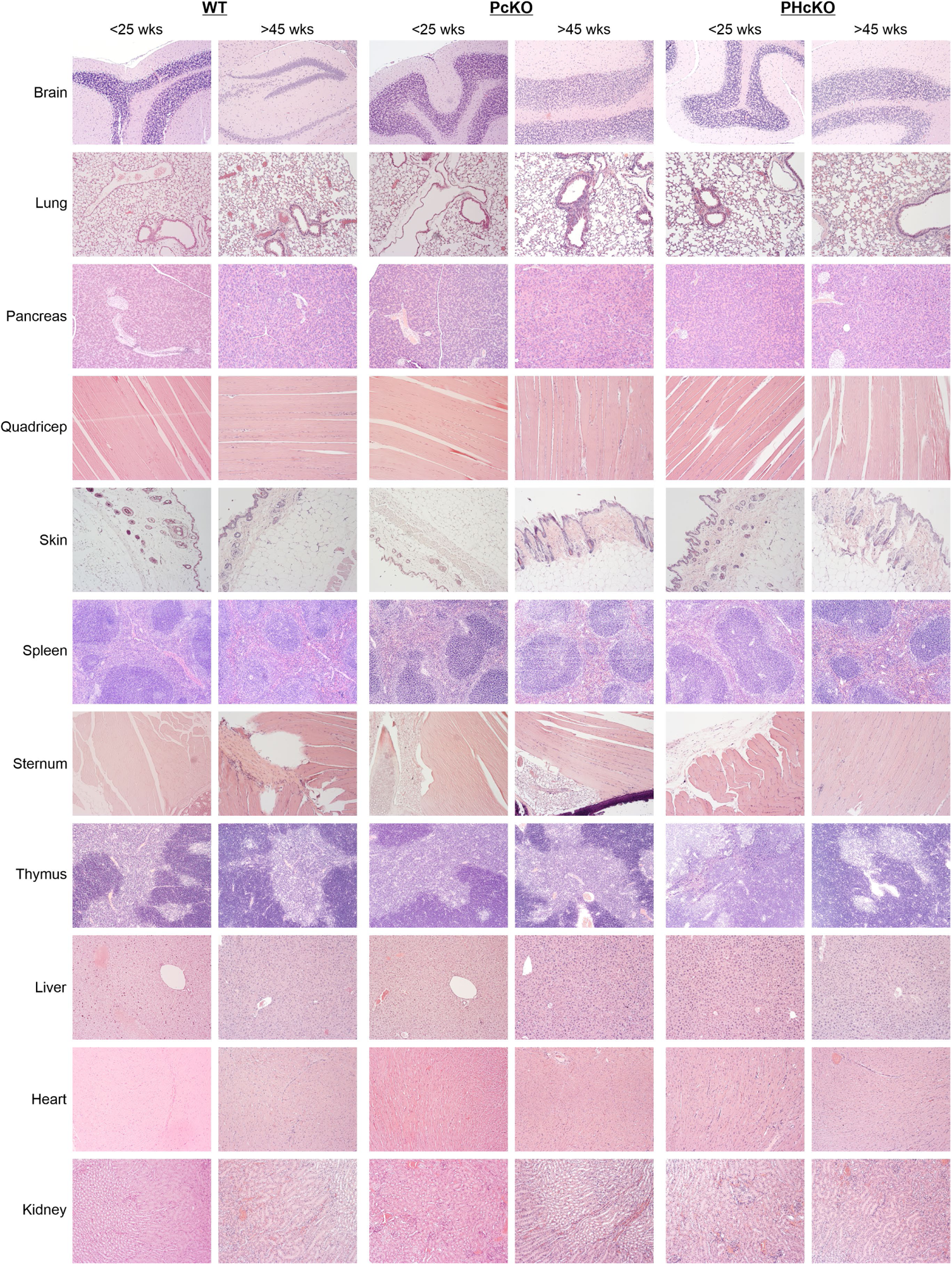
Tissue histology of WT, PcKO, and PHcKO animals was unchanged. H&E stained tissue sections from all three genotypes and across two age groups as indicated.

In summary, the blood cellularity and histologic analyses demonstrate a lack of overt phenotype in both the PcKO and PHcKO mouse strains in both young and old age groups.

### Platelets do not impact plasma glycoprotein glycans

Most glycoproteins in plasma are synthesized and released from hepatocytes, with the notable exception of IgG and other antibodies. Using the HcKO mouse, we previously showed that the lack of ST6Gal1 in hepatocytes generated robust differences in overall plasma glycoprotein glycosylation (Oswald, D.M., Jones, M.B., et al. 2020). This included the expected loss of α2,6-linked sialylation as detected by the lectin SNA, but also a reduction in α1,6-core fucosylation as measured by LCA and an increase in bisecting GlcNAc residues as measured by PHA-E. With an ELISA-like plate assay using lectins to detect glycan composition, we characterized total plasma glycoprotein glycosylation (Fig 3). We found that none of the lectin-detected glycans were altered in the PcKO mouse compared to WT controls. However, we found a dramatic loss of SNA staining in PHcKO animals compared to WT animals, and this change was mirrored in HcKO controls. Similarly, LCA staining was decreased and GlcNAc-detecting WGA was increased in both PHcKO and HcKO plasma. The PHA-E signal was increased in HcKO plasma as found previously (Oswald, D.M., Jones, M.B., et al. 2020), but despite the upward trend in PHcKO plasma, it did not reach statistical significance when compared to WT. Overall, the changes in plasma glycoprotein glycosylation are driven solely by the loss of ST6Gal1 in hepatocytes and not in platelets.

**Figure 3.**
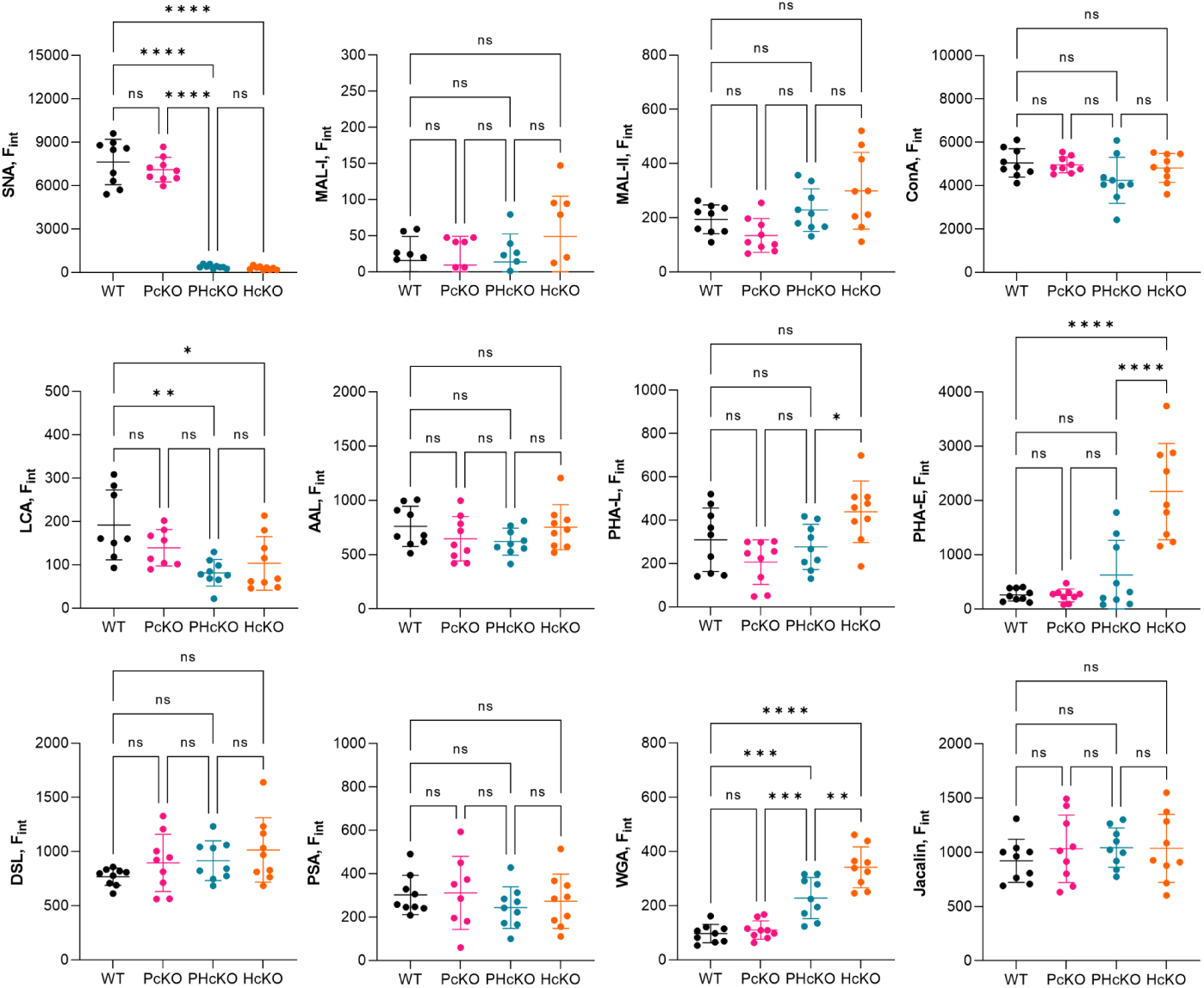
Plasma glycoprotein glycans were altered in PHcKO but not PcKO animals. Lectin-based ELISA-like plate assays of plasma from WT, PcKO, PHcKO, and HcKO animals using a panel of lectins, including SNA for α2,6-linked sialic acids, LCA for α1,6 core fucosylation, PHA-E for bisecting GlcNAc. *p < 0.05; **p < 0.01; ***p < 0.001; ****p < 0.0001; N=9

### PcKO and PHcKO animals have differential tissue sialylation

In order to characterize any change in tissue sialylation resulting from both genetic changes in the PcKO and PHcKO animals, we stained a range of tissues from young mice with fluorescent SNA, collected images using confocal microscopy (Fig 4A), and quantified the mean fluorescence to quantitatively compare between strains (Fig 4B). First, PcKO tissues showed a minor reduction in α2,6-sialylation in the liver and modest increases in both the heart and spleen. In all of these cases, the difference is only apparent as an average across multiple animals (Fig 4B). An increase in α2,6-sialylation was seen in the lungs of PcKO animals compared to their WT counterparts, while no differences were found in the kidney, gut, pancreas, brain, or thymus. Second, the PHcKO tissues showed greater change compared to PcKO animals (Fig 4). As expected, the liver was dramatically reduced in SNA staining, but SNA signal was also significantly reduced in the spleen, lung, and kidney of PHcKO mice. The modest increase in heart sialylation was the same as found in PcKO animals, while PHcKO animals showed a substantial increase in SNA staining in the brain. As seen with the PcKO tissues, no change in the gut, pancreas, or thymus was apparent in the PHcKO mice.

**Figure 4.**
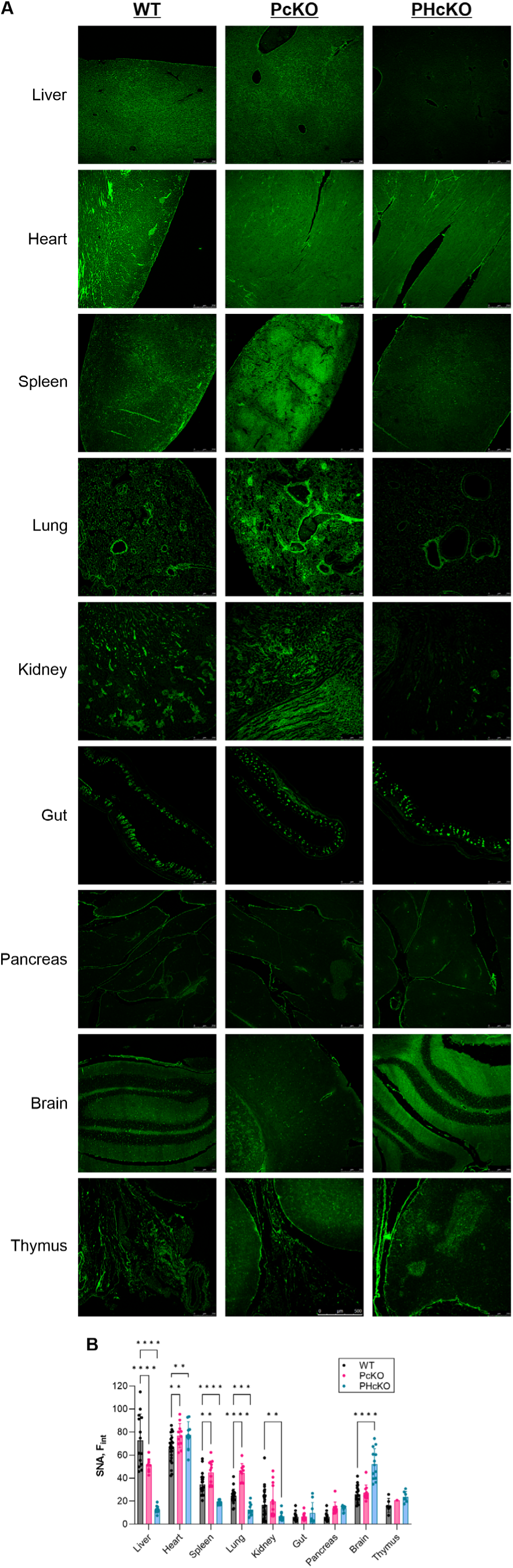
SNA staining of tissues by confocal microscopy revealed changes in tissue sialylation. (A) Representative images from each tissue and animal strain. All mice were in the young age group (<20 wks of age). (B) Quantitation of SNA signal intensity across at least 3 animals and many images. *p < 0.05; **p < 0.01; ***p < 0.001; ****p < 0.0001; N=6-30

### Platelet ST6Gal1 does not impact EAE susceptibility

We have already established that loss of ST6Gal1 in hepatocytes translates into greater susceptibility to T cell-mediated autoimmunity, including EAE (experimental autoimmune encephalomyelitis), the murine model of multiple sclerosis (Oswald, D.M., Zhou, J.Y., et al. 2020). Moreover, the depletion of platelets has been proposed to ameliorate the effects of EAE, suggesting a connection between platelets and autoimmunity (Langer, H.F., Choi, E.Y., et al. 2012). We therefore sought to determine the impact of ST6Gal1 ablation in platelets on EAE pathogenesis. We induced EAE in PcKO and WT mice using a standard immunization protocol of myelin oligodendrocyte glycoprotein (MOG) peptide and scored them for disease and weight loss over a time course of 28 days. PcKO mice developed disease slightly faster, with a change in disease score and weight becoming apparent two days earlier than in WT controls (Fig 5A-B). However, we did not observe a change in overall disease severity as measured by weight loss or disease score, even observing the same rebound in weight that occurred approximately 20 days post-EAE induction in WT mice. Tissue sections of the brain taken on day 28 failed to reveal differential disease as assessed by cellular infiltration (H&E; Fig 5C) or the degree of myelin loss (Luxol Fast Blue; Fig 5D). These findings suggest that the knockout of ST6Gal1 from the platelets of mice does not impact the severity of a model of T cell-dependent autoimmunity.

**Figure 5.**
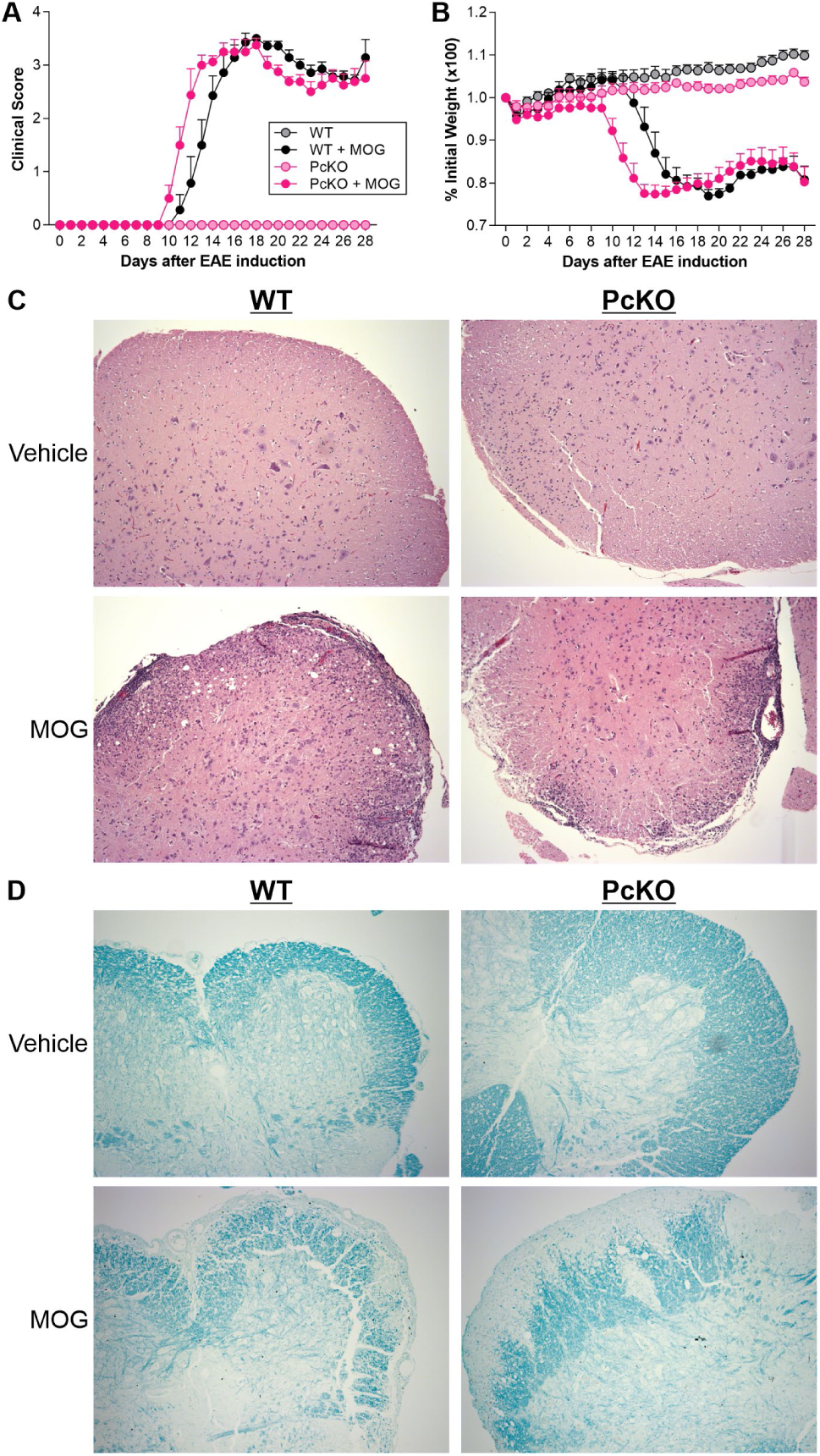
PcKO animals have similar susceptibility to EAE. Animals were placed on the EAE protocol with MOG peptide or vehicle control and monitored by clinical score (A) and weight (B) on a daily basis. At day 28, mice were sacrificed and brains were analyzed for pathology using both H&E (C) and Luxol Fast Blue (D) staining to show infiltrating immune cells and myelin respectively.

### Platelet ST6Gal1 does not change IgG sialylation

Overall, the characterization of the PcKO and PHcKO animals revealed mostly predicted changes in glycosylation, but no spontaneous pathology nor obvious change in susceptibility to autoimmunity was found. Since the goal of creating these animals was to determine the potential role for platelets and the granular microenvironment in mediating IgG sialylation, we compared the amount of α2,6 sialylation on purified IgG from WT, PcKO, PHcKO, and HcKO mice using lectin staining. We found no difference in SNA staining between any of the strains (Fig 6A). We further analyzed both WT and PcKO IgG using hydrophilic interaction liquid chromatography (HILIC). In agreement with the lectin staining assay, we found no difference in the number of glycans containing zero, one, or two sialic acid residues (Fig 6B).

**Figure 6.**
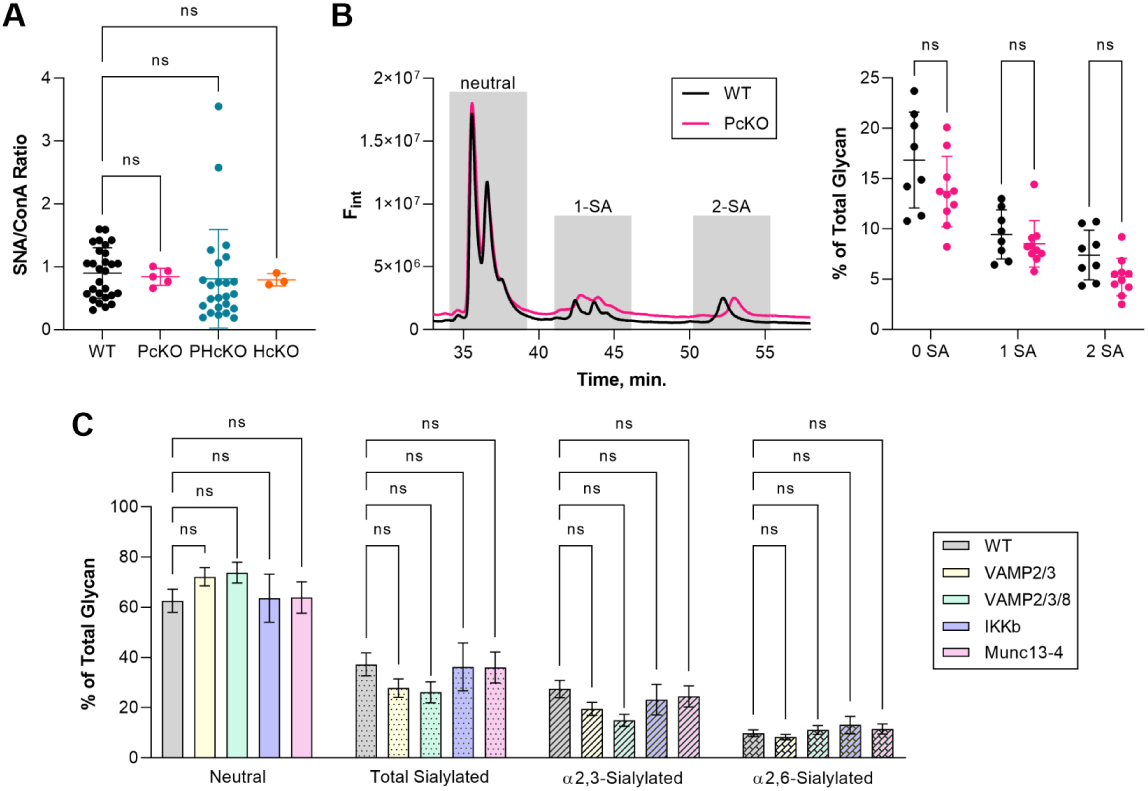
IgG sialylation remains unchanged in PcKO, PHcKO, and platelet granule release mutant animals. (A) Lectin-based ELISA-like plate assay of purified IgG and stained with SNA and ConA. The SNA/ConA ratio is shown for comparison. N=3-28. (B) HILIC HPLC analysis of purified IgG from WT and PcKO mice. Quantitation of the neutral, singly sialylated, and dually sialylated species is shown on the right with representative HILIC profiles on the left. N=8-10. (C) Mass spectrometry analysis of N-glycans of purified IgG from the indicated platelet granule release mutants. N=2 samples per analysis. ns = not significant.

Soluble N-ethylmaleimide sensitive factor attachment protein receptors (SNAREs) are a family of proteins responsible for platelet exocytosis, including the release of dense and α-granules. These are also known as vesicle-associated membrane proteins, or VAMPs, and VAMP2, VAMP3, VAMP7, and VAMP8 are those most relevant for mouse platelets (Joshi, S., Banerjee, M., et al. 2018). Defects in these molecules and others involved with their regulation contributes to reductions in granule release, which should reduce IgG sialylation within the circulation if platelets are a key to IgG remodeling. We compared the IgG sialylation from several mouse strains with exocytosis defects, including a Pf4-specific VAMP2/3 double and VAMP2/3/8 triple knockout (Joshi, S., Banerjee, M., et al. 2018), IKKβ knockout (Karim, Z.A., Zhang, J., et al. 2013), and the Munc13-4 knockout (Ren, Q., Wimmer, C., et al. 2010). Purified IgG samples were analyzed using glycan mass spectrometry as before (Oswald, D.M., Lehoux, S.D., et al. 2022). Consistent with the results from PcKO and PHcKO animals, we found no change in total sialylation or α2,6-specific sialylation on these IgG molecules (Fig 6C).

## Discussion

We have previously demonstrated that IgG sialylation is dependent on ST6Gal1, as shown by the total loss of α2,6-sialylation on IgG in a global ST6Gal1 knockout (Oswald, D.M., Lehoux, S.D., et al. 2022). Our previous studies have also shown that ST6Gal1 in B cells is dispensable for sialylating IgG, suggesting that IgG sialylation occurs in a B cell-extrinsic manner (Jones, M.B., Oswald, D.M., et al. 2016). We further demonstrated that the failure of B cells to robustly sialylate IgG is primarily due to intracellular trafficking of nascent IgG in antibody-secreting cells such that contact between ST6Gal1 and IgG is limited (Glendenning, L.M., Zhou, J.Y., et al. 2022). These findings indicate that IgG sialylation occurs through a currently unidentified pathway.

The first candidate site for IgG sialylation was the plasma itself, as ST6Gal1 is known to be secreted from the liver upon proteolytic release from the membrane by BACE1 (Kitazume, S., Oka, R., et al. 2009, Woodard-Grice, A.V., McBrayer, A.C., et al. 2008). In fact, studies using a mouse with a reported liver-specific loss of ST6Gal1 through the ablation of the P1 promoter sequence upstream of ST6Gal1 (Appenheimer, M.M., Huang, R.Y., et al. 2003) showed a reduction in IgG sialylation (Jones, M.B., Nasirikenari, M., et al. 2012). Importantly, we and others have found that circulatory ST6Gal1 retains its activity (Jones, M.B., Oswald, D.M., et al. 2016), supporting the hypothesis that circulatory ST6Gal1 may be responsible for IgG sialylation. We therefore created a hepatocyte-specific knockout of ST6Gal1 (HcKO). While this mouse has no detectable ST6Gal1 in the circulation, the degree of IgG sialylation was unchanged (Oswald, D.M., Lehoux, S.D., et al. 2022). We concluded that IgG sialylation is primarily not occurring within the plasma.

The next candidate site for IgG sialylation was platelets. We hypothesized that IgG sialylation occurs within platelet granules because they have been shown to contain all of the necessary molecules – IgG, CMP-sialic acid, and ST6Gal1. Yet, it is not entirely clear whether the ST6Gal1 present in the platelets are endogenously produced within the MK/platelets themselves or endocytosed from the circulatory environment. Therefore, we created two novel mouse strains, one lacking ST6Gal1 in the MK/platelet population (PcKO), and another lacking ST6Gal1 in both the MK/platelet population and the circulatory environment (PHcKO). Once again, however, we report that IgG sialylation was unchanged. This suggests that although IgG sialylation is occurring in a B-cell extrinsic manner, it is not dependent on the activity of plasma-localized ST6Gal1, does not require ST6Gal1 expression in platelets, and does not require platelet endocytosis of hepatocyte-released ST6Gal1 from the plasma.

Although we demonstrate that platelet-localized ST6Gal1 does not make a significant impact on IgG sialylation, these data do not rule out the possibility that platelets could play some role in the pathway through the release of CMP-SA or IgG sequestration. Thus, there are two major questions that arise from these data. Firstly, if all of the components required for IgG sialylation are present within platelets, why does it not occur? Based on our previous work, we suspect that these components may not share the same intracellular compartment. It may be true that IgG, ST6Gal1, and CMP-SA are all found in platelet granules, but it is unclear whether they are coming into contact within the same granule and at a high enough concentration to efficiently promote IgG sialylation. Reports indicate that IgG is present in α-granules (George, J.N. 1991, Verheul, H.M., Lolkema, M.P., et al. 2007) and that ST6Gal1 and CMP-SA are released by activated platelets through degranulation (Jones, M.B., Oswald, D.M., et al. 2016, Wandall, H.H., Rumjantseva, V., et al. 2012), but a direct co-localization between IgG and ST6Gal1 has not been demonstrated. In light of our findings, we therefore hypothesize that platelets fail to make a significant impact on IgG sialylation due to a lack of direct IgG/ST6Gal1/CMP-SA contact within the platelet, exactly as we have recently reported with antibody-secreting B cells which fail to co-localize ST6Gal1 and IgG (Glendenning, L.M., Zhou, J.Y., et al. 2022).

Second, if not in B cells, the plasma, or platelet granules, where does IgG sialylation occur? There are no published findings that identify where IgG is sialylated *in vivo*, yet localizing the pathway responsible for IgG sialylation is a prerequisite for dissecting the regulatory elements which impact glycan-sensitive IgG function. This applies to endogenously produced IgG as well as exogenously provided IgG and Fc-fusion drug delivery constructs. When sialylation of endogenous IgG is driven by intravenous injection of the relevant enzymes and CMP-sialic acid, murine rheumatoid arthritis can be ameliorated (Pagan, J.D., Kitaoka, M., et al. 2018). This shows that manipulation of the IgG sialylation regulatory pathway, wherever it may be, could prove a valuable target for treating autoimmune and inflammatory diseases through changing endogenous IgG activity. It is also critical to understand how the regulation of this pathway may impact exogenous antibody therapies. Recent studies have shown that B-cell extrinsic sialylation of Fc domains of exogenous antibody can occur, although changes to the glycans on the Fab region are observed first due to increased accessibility of this glycan (Schaffert, A., Hanic, M., et al. 2019).

Overall, it is clear that endogenous IgG glycans are remodeled *in vivo* after release from antibody-secreting cells, and that the same remodeling pathway is likely able to manipulate exogenous IgG, affecting its overall immune function. The regulatory mechanisms underlying homeostatic IgG sialylation may therefore provide a novel group of targets to be developed as glycan-based therapeutics in the future. But, until the pathway for IgG sialylation is found, understanding the underlying regulation of IgG glycan remodeling will remain a significant knowledge gap in the human immune system.

## Experimental Procedures

### Mice

All animal work was approved by the Institutional Animal Care and Use Committee (IACUC) of Case Western Reserve University. Mice were procured from Jackson Labs and were housed and bred in the CWRU Animal Resource Center in accordance with IACUC guidelines. ST6Gal1-flox mice (Jackson Labs, stock 006901) and Alb-Cre mice (Jackson Labs, stock 003574) were crossed to produce the HcKO line, as we document in our previous manuscripts (Oswald, D.M., Jones, M.B., et al. 2020, Oswald, D.M., Zhou, J.Y., et al. 2020). Wherever the phrases “wild type” or “parental” are used, they refer to mice of the ST6Gal1f/f background, which carry no phenotype nor change in ST6Gal1 expression. PcKO mice were produced by crossing the ST6Gal1-flox mice with Pf4-Cre mice (Jackson Labs, stock 008535). PHcKO mice were created by crossing HcKO and PcKO mice, resulting in mice carrying ST6Gal1-flox, Alb-Cre, and Pf4-Cre alleles.

### Flow cytometry

Flow cytometry was performed on bone marrow-derived cells, which were isolated by exposing the bone marrow from murine femurs and tibias, and spinning the marrow out at 2000×g for 10 min. Bone marrow from each mouse was then combined and subjected to red blood cell lysis in 1X BD PharmLyse (BD Biosciences, 555899). Cells were then washed with PBS, filtered through a 70 μm nylon filter, and blocked for 30 min in carbohydrate-free blocking solution (Vector Labs) and Fc block. Cells were stained with SNA-FITC (Vector Labs) and Pf4-APC (Abcam, ab280969). Flow cytometry was run on a BD Symphony S6 and data were analyzed using FlowJo.

### Tissue sectioning, staining, and imaging

All harvested tissues were fixed in 10% formalin (VWR, Radnor, PA) for 24 hr and sent to AML Laboratories (Jacksonville, FL) for paraffin embedding and sectioning. Sectioned slides were stained with H&E or Luxol Fast Blue. Images were acquired using a High-Speed Microscope Camera (Amscope, Irvine, CA). For confocal imaging, tissue sections were stained with fluorescent SNA for 1 hour at room temperature, washed, and mounted to slides for image acquisition on a Leica TCS SP5 confocal microscope (Leica, Wetzlar, Germany). Image quantitation by fluorescence intensity was performed using ImageJ.

### Blood cellularity analysis

Whole blood was isolated from mice through a cardiac puncture immediately following sacrifice via CO_2_ inhalation. Cellularity was analyzed on a Heska HemaTrue Veterinary Hematology Analyzer.

### EAE model of multiple sclerosis

Age-matched female mice between 11 and 23 weeks underwent EAE induction using the MOG35-55/ CFA Emulsion and pertussis toxin (PTX) kit from Hooke Laboratories (Lawrence, MA) according to manufacturer’s instructions. Mice were immunized with 200 mg of MOG35-55/CFA emulsion subcutaneously in two locations in the back on day 0 and were administered 100 ng of PTX intraperitoneally on days 0 and 1 of the trial. Negative controls for disease received a CFA emulsion and PTX according to the same schedule. Mice were weighed and scored daily according to Hooke Laboratory’s EAE scoring guide (https://hookelabs.com/protocols/eaeAI_C57BL6.html). Mice were euthanized if they reached a disease score of 5 according to IACUC animal welfare guidelines.

### IgG purification

IgG was purified from plasma samples by separation over a HiTrap Protein A HP Antibody Purification Column (GE Life Sciences) fitted to a GE Life Sciences Akta Purifier 10 HPLC. Binding and washing were done with 10 mM Tris, pH 7.5, and 150 mM NaCl. Elution of IgG was done with 50 mM Citrate, pH 4.5, and 150 mM NaCl. Purified IgG was buffered exchanged into PBS for ELISA and long-term storage, or water for HPLC glycan analyses. Purity validated by SDS-PAGE and Coomassie blue staining (not shown).

### Glycan HPLC analysis

Hydrophilic interaction liquid chromatography (HILIC) analysis of intact N-glycans was performed using 2-aminobenzamide (2-AB) labeling and separation on a Zorbax NH2 column (Agilent), essentially as described elsewhere (Bigge, J.C., Patel, T.P., et al. 1995, Guile, G.R., Rudd, P.M., et al. 1996). IgG was digested with trypsin (ThermoFisher) overnight at 37°C. Trypsin was inactivated by heating to 95°C for 10 min. N-glycans were enzymatically removed from the tryptic peptides with PNGase F (NEB) overnight at 37°C. Glycans were separated from peptides by reversed-phase chromatography through a C18 cartridge (ThermoFisher) and dried in a lyophilizer (LabConco) overnight. Purified glycan was labeled using 50 μg/μl 2-AB and 60 μg/μl sodium cyanoborohydride in a 3:7 solution of acetic acid and DMSO at 65°C for 3 h. Unreacted label was removed on a G-10 desalting column (GE Healthcare). HILIC analysis was performed using a 36–45% gradient of 100 mM ammonium formate, pH 4.4, and acetonitrile. 2-AB fluorescence was detected with an excitation of 330 nm and emission of 420 nm.

### Lectin ELISA

Lectin ELISA was performed on purified IgG as published previously with plasma samples (Oswald, D.M., Sim, E.S., et al. 2019). Briefly, purified IgG was diluted to 1 mg/ml in PBS, pipetted into a 96-well high-binding ELISA plate (Microlon High Binding; Greiner BioOne), and incubated overnight at 4°C. The plate was blocked with carbohydrate-free blocking solution (Vector Labs) for 1 h at room temperature. Biotinylated lectins (Vector Labs) were diluted to 1 μg/ml in carbohydrate-free blocking solution and incubated on the plate for 1 h at room temperature. Signal was detected using europium-conjugated streptavidin (Perkin Elmer) and time-resolved fluorescence as measured in a Victor Nivo multilabel plate reader.

### Mass spectrometry

Mass spectrometry N-glycan profile and sialic acid linkage analysis was performed on purified IgG exactly as previously reported (Alley, W.R., Jr. and Novotny, M.V. 2010, Oswald, D.M., Lehoux, S.D., et al. 2022). Briefly, IgG was digested with Trypsin (Millipore-Sigma) overnight, followed by PNGaseF (New England Biolabs) release of N-glycans for 20 hours. The PNGaseF-released N-glycans were isolated on a C18 Sep-Pak column and evenly split into two samples.

One of the two dried N-glycan aliquots for each sample was permethylated using standard methods (Oswald, D.M., Lehoux, S.D., et al. 2022), cleaned on a C18 Sep-Pak column, and used for standard MS analysis (below). For the DMT-MM (4-(4,6-dimethoxy-1,3,5-triazin-2yl)-4-methylmorpholinium chloride) treatment to differentiate α2,3 and α2,6 sialic acid linkages, a solution of 0.5 M of DMT-MM (Millipore-Sigma) was prepared in 500 mM of NH_4_Cl, pH 6.5. 10 µL of the DMT-MM solution were added to the second aliquot of each N-glycan sample and incubated at 60°C for 15 hours. Samples were cleaned on a C18 Sep-Pak column, lyophilized, and permethylated as before.

Permethylated N-glycans with and without DMT-MM derivatization were spotted on a MALDI polished steel target plate (Bruker Daltonics). MS data were acquired on a Bruker UltraFlex II MALDI-TOF Mass Spectrometer instrument. The reflective positive mode was used, and data were recorded between 500 and 6000 m/z. For each MS N-glycan profile, the aggregation of 20,000 laser shots or more were considered for data extraction. Only MS signals matching an N-glycan composition were considered for further analysis. Subsequent MS post-data acquisition analysis was made using mMass (Strohalm, M., Kavan, D., et al. 2010).

### Data and statistical analysis

All data were analyzed and plotted using GraphPad Prism. Statistical analyses used were Student’s t-test or one-way ANOVA where appropriate (*p < 0.05; **p < 0.01; ***p < 0.001; ****p < 0.0001)

## Acknowledgements

The authors wish to acknowledge thoughtful discussion and experimental input from Douglas Oswald, Jill Cavanaugh, and Lori Kreisman during the completion of this study.

## Author Contributions

LMG, experimental design, data collection and analysis, manuscript writing; JYZ and ENK, data acquisition; CG and RDC, mass spectrometry data collection and analysis; SJ and SWW, sample acquisition from murine models of platelet granule release; BAC, experimental design, data collection and analysis, manuscript writing, funding.

## Abbreviations

4-(4,6-dimethoxy-1,3,5-triazin-2yl)-4-methylmorpholinium chloride: (DMT-MM)
cytidine monophosphate N-acetylneuraminic acid: (CMP-sialic acid)
hepatocyte-specific conditional knockout of ST6Gal1: (HcKO)
megakaryocyte: (MK)
MK/platelet-specific conditional knockout of ST6Gal1: (PcKO)
MK/platelet- and hepatocyte-dual specific conditional knockout of ST6Gal1: (PHcKO)
Platelet factor 4: (Pf4)
rheumatoid arthritis: (RA)
myelin oligodendrocyte glycoprotein: (MOG)

